# Complete genomes of *Rickettsia typhi* reveal a clonal population

**DOI:** 10.1101/2025.05.25.654783

**Authors:** Chantisa Keeratipusana, Weerawat Phuklia, Vanheuang Phommadeechack, Janjira Thaipadungpanit, Vilada Chansamouth, Koukeo Phommasone, Sayaphet Rattanavong, Catrin E. Moore, Matthew T. Robinson, Allen L. Richards, Paul N. Newton, Elizabeth M. Batty

## Abstract

Murine typhus, caused by infection with *Rickettsia typhi*, is a neglected disease contributing to infectious disease burden in south- and southeast Asia. Despite its importance, we have minimal knowledge of the genomics of *R. typhi*, with only four complete genomes being sequenced prior to this work. We sequenced a further 25 genomes including historical strains collected before 1976 from both human and rat hosts, and recent genomes isolated from patients at a single hospital in Laos. Whole genome SNP analysis reveals extremely low levels of genetic diversity across the 29 genomes, with overall nucleotide diversity (π) of 1.27e^−05^ and evidence of purifying selection, and a minimal pan-genome. Phylogenetic analysis shows clustering of the genome by historic or modern origin, with the exception of one modern strain which is most closely related to historic strains from Thailand, and no clustering by host origin. The highly conserved genome of *R. typhi* suggests strong constraints on genome evolution in this obligate intracellular parasite, and has implications for the design of future murine typhus diagnostic tools and vaccines.

**Impact Statement:** *Rickettsia typhi* is the causative agent of murine typhus, an infectious disease that causes morbidity and mortality in humans. This study has expanded the genomic dataset from a small number of genomes to a comprehensive collection of 29 complete genomes. The genome comparison includes historic and modern strains and reveals a clonal population structure with minimal genetic diversity across time. This shows the evolutionary stability of *R. typhi*, which is most likely due to strong purifying selection and an intracellular lifestyle. This extremely low genomic variation has significant impacts for the development of murine typhus vaccines and robust diagnostic assays.

## Introduction

*Rickettsia typhi* causes murine typhus, a neglected infectious disease contributing to the burden of febrile illness and associated morbidity and mortality in humans, especially in south- and southeast Asia [1]. It is transmitted by the bites of infected fleas, which typically live on rats [2]. Murine typhus occurs worldwide and while mortality is low at approximately 0.4-3.6% [3], morbidity in untreated disease can be significant, showing the importance of early diagnosis and treatment.

All species of the genus *Rickettsia* are obligate intracellular bacteria associated with arthropod hosts, and can be classified into four major groups: 1) the typhus group, which includes *R. typhi* and *Rickettsia prowazekii*, that cause murine typhus and epidemic typhus, with flea and louse hosts, respectively; 2) the spotted fever group, which includes *Rickettsia rickettsii, Rickettsia japonica* and *Rickettsia conorii*, found in ticks; 3) the ancestral group of *Rickettsia bellii* and *Rickettsia canadensis*, which are non-virulent species found in ticks; and 4) the transitional group rickettsiae which include: *Rickettsia akari, Rickettsia australis* and *Rickettsia felis*, which are mite-, tick- and flea-borne, respectively [4].

The first complete genome of *R. typhi* was sequenced in 2004, revealing 877 genes in a 1.11Mb genome [5]. Most *Rickettsia* species contain a single rRNA operon composed of the 16S, 23S, and 5S rRNA genes. This configuration is consistent with their relatively low replication rates and adaptability to a stable intracellular environment. *Rickettsia* species share a single rRNA operon, with little variation [6]. Further comparative genomics of complete rickettsial genomes defined five phylogenetic groups, with the typhus group having the smallest genome size of the five groups. While plasmids were identified in many rickettsial species, no plasmids have been identified in any of the typhus group strains, although *R. prowazekii* has a gene cluster thought to result from integration of an ancestral *Rickettsia* plasmid into the chromosome [7]. *R. typhi* is naturally sensitive to rifampin, but other *Rickettsia* species in the spotted fever group show resistance, and a limited number of studies have investigated laboratory-derived rifampin resistance in *R. typhi* [8–10].

While the first genome was published over twenty years ago, our knowledge of *R. typhi* genomics remains minimal, likely due to low rates of diagnosis and the difficulty in culturing and sequencing bacteria from patients even when a confirmed diagnosis is available [11]. Previous work has compared four complete genomes and identified a small number of genetic changes between strains collected over a long timeframe [12]. The mutations detected in three of these genomes were used as probes to type a further 15 isolates of *R. typhi*, and to generate a phylogeny based on 26 SNPs and indels [13].

In this study we used whole genome sequencing to expand our knowledge on the genomes of *R. typhi* from two populations: a set of eleven historic strains collected from seven countries between 1928 and 1975, from the collection of the Naval Medical Research Center, USA, and a set of seventeen samples collected over ten years (2006-2019) from Mahosot Hospital in the Lao PDR (Laos) [11]. Using these datasets we were able to compare historic data to modern strains, and describe the diversity found in patient isolates collected over a ten-year timeframe in one country.

## Materials and Methods

### *Rickettsia typhi* culture and DNA extraction

For the samples cultured from patients with murine typhus at Mahosot Hospital, up to 5 ml of blood was drawn into EDTA tubes after obtaining written informed consent from patients. Full isolation procedure is detailed in Ming *et. al*. [11]. Briefly, either 1 mL whole EDTA anticoagulated blood, or 200 µL EDTA buffy coat fraction was mixed with 5 mL of RPMI 1640 medium supplemented with 10% foetal calf serum (FCS) (PAA Laboratories, USA, or Gibco, USA). Inoculation was performed directly in 12.5cm^2^ flasks containing L929 (ATCC^®^ number CCL-1^™^) cell monolayers at 80% confluence, and subsequently centrifuged for 30 minutes at 50 ×*g*. Flasks were incubated for 2 hours at 35°C in a 5% CO_2_ atmosphere and then the inoculum washed and replaced with RPMI 1640 with 10% FCS. Flasks were incubated for a minimum of 8 weeks at 35°C in a 5% CO_2_ atmosphere, with a twice-weekly media change and sub-culture performed at week four. Cells were checked for successful isolation by indirect IFA using pooled antibodies against scrub typhus, spotted fever and typhus group rickettsial agents every week after 4 weeks of incubation or earlier [14]. Cultures positive by IFA underwent DNA extraction using DNeasy Blood & Tissue Kit (Qiagen, UK) and isolation was confirmed by qPCR targeting the *R. typhi* 17kDa gene [15].

For genomic DNA (gDNA) extraction, infected cells from three 75-cm^2^ flasks were mechanically detached, resuspended and transferred to a 50-mL conical-bottom tube and centrifuged at 3,220 x g for 10 minutes. The pellet was resuspended in 3-mL fresh medium and transferred to 1.5 mL microcentrifuged tubes. Tubes were vortexed for one minute and host cell debris was removed after centrifugation at 300 x g for 3 minutes. The supernatant was passed through a µM filter (Paradisc, GE Whatman, USA). Subsequently, 10 µL of 1.4 µg/µL DNAse was added per 1 mL of bacteria solution, incubated at room temperature for 30 minutes. The mixture was centrifuged at 18,188 x g for 10 minutes, the pellet was washed twice with 0.3M sucrose (Sigma-Aldrich). DNA was extracted from the *R. typhi* pellet using the DNeasy Blood & Tissue kit (Qiagen, Hilden, Germany) following the manufacturer’s protocol. DNA was eluted in AE buffer and stored immediately at −20°C. The yield of genomic DNA was quantified using a nanodrop spectrophotometer (Nanodrop 2000, Thermo Scientific, UK). The ratio between *R. typhi* and host cells was determined using qPCR of the *R. typhi ompB* [16] and mouse *GADPH* genes.

### Illumina sequencing

Sequencing libraries were prepared from extracted DNA using the Illumina Nextera Flex library preparation kit (Illumina, USA) according to manufacturer’s protocols. Libraries were barcoded using the Illumina DNA Tagmentation indexes (Illumina, USA) and pooled and sequenced on a single run of a MiSeq instrument using the MiSeq V3 paired-end 300bp sequencing kit.

### Bioinformatic analysis

Three public assembly genomes and 25 Illumina short reads were analysed using the Bactopia v2.2.0 pipeline [17]. Variants were called and annotated using freeBayes v1.3.6 [18] and snpEff v5.2 [19], respectively. Consensus data with only substitution variants were used to create a phylogenetic tree with 1000 bootstrap replicates using Iqtree v2.2.5 [20]. The tree was plotted, annotated and visualized using ggtree v3.10.0 [21]. BactDating [22] was used to assess the evolutionary rate, and SNPgenie [23] to estimate nucleotide diversity and selection pressure. Illumina short reads were *de novo* assembled using Skesa [24]. After *de novo* assembly, contigs showing similarity to the mouse genome with a 99% identity threshold match using BLAST v2.15.0 [25] were removed. BUSCO v5.7.1 [26] using the OrthoDB version10 bacterial dataset was used to assess the gene content. Prokka v1.14.6 [27] was used to annotate both the existing *R. typhi* genomes and the new assemblies. Panaroo v1.3.4 [28] was used to construct a pan-genome with strict stringency mode and core-genome sample threshold of 1. TM2540 assembly data was improved by TM2540 Illumina sequencing data in this study using Pilon v1.24 [29]. The polished TM2540 assembly data was compared to the original TM2540 by the dnadiff option from MUMmer v3.23 [30].

### Ethics approval

Ethical approvals for collection of samples for determination of the etiology of fever in Lao were granted by the Lao National Ethics Committee for Health Research and the Oxford Tropical Research Ethics Committee.

### Data availability

The three existing *R. typhi* genomes were downloaded from RefSeq under the accession numbers NC_006142.1 (Wilmington), NC_017066.1 (TH1527), and NC_017062.1 (B9991CWPP). The sequence data from this project are available in the NCBI Sequence Read Archive under project PRJNA1127778.

## Results

We generated whole genome sequencing data from two populations of *R. typhi*: a set of eleven historic strains collected from seven countries between 1928 and 1975, and a set of fourteen strains collected over ten years (2006-2019) from a single hospital in Laos (Table 1). This second set included resequencing of the previously sequenced TM2540 isolate [12]. We included three complete genomes that were previously made available in our analysis (TH1527, B9991CWPP, and Wilmington/NC_006142.1), and the Wilmington strain was used as the reference genome. The date and country of origin for each strain is shown in Table 1. Whole genome sequencing statistics are shown in Table 2. Coverage depth ranged from 37X to 294X. 1%-11% of reads were determined to be mouse contamination, which was expected as the *R. typhi* strains were cultured in L929 mouse fibroblast cells.

**Table 1.**
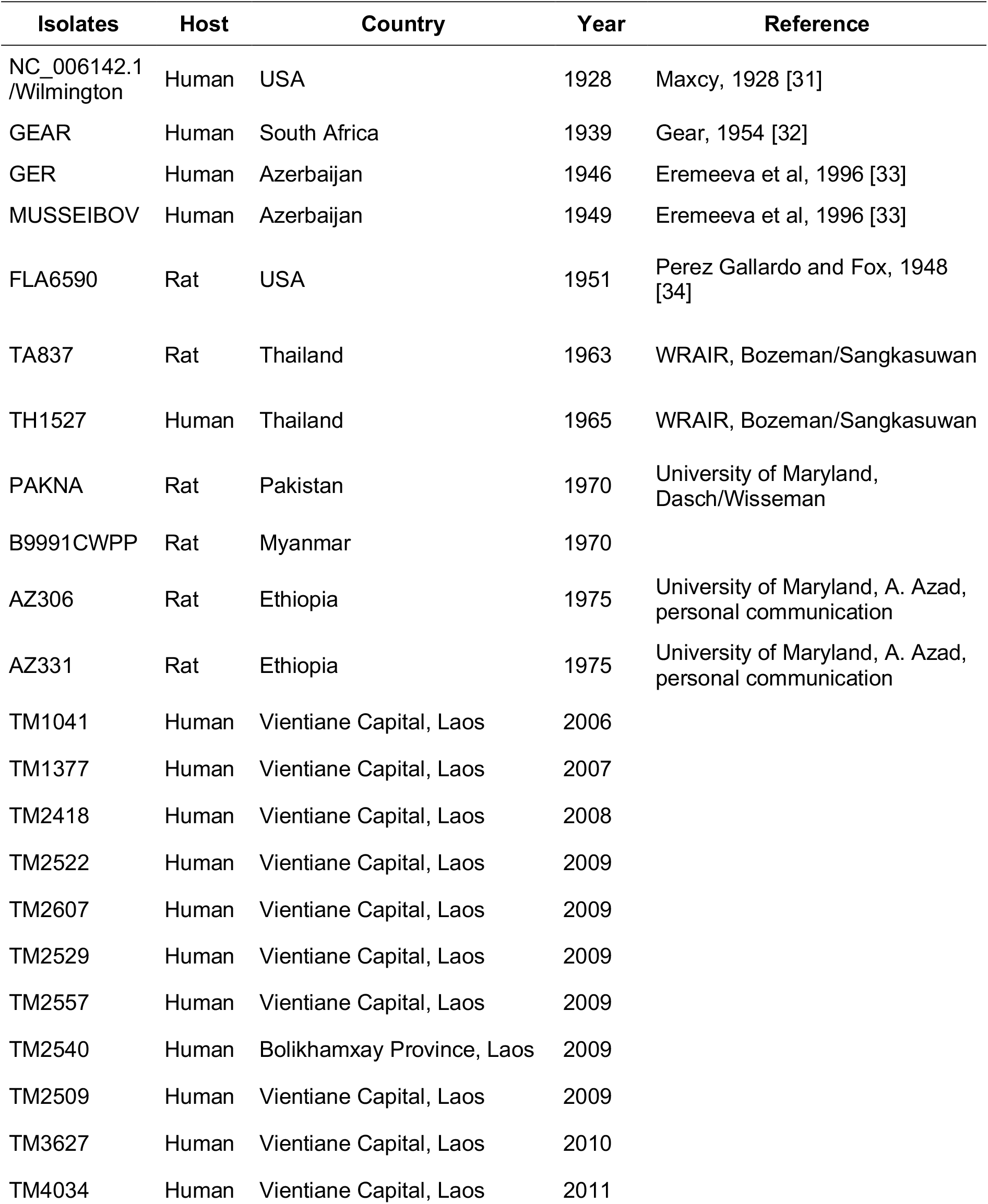
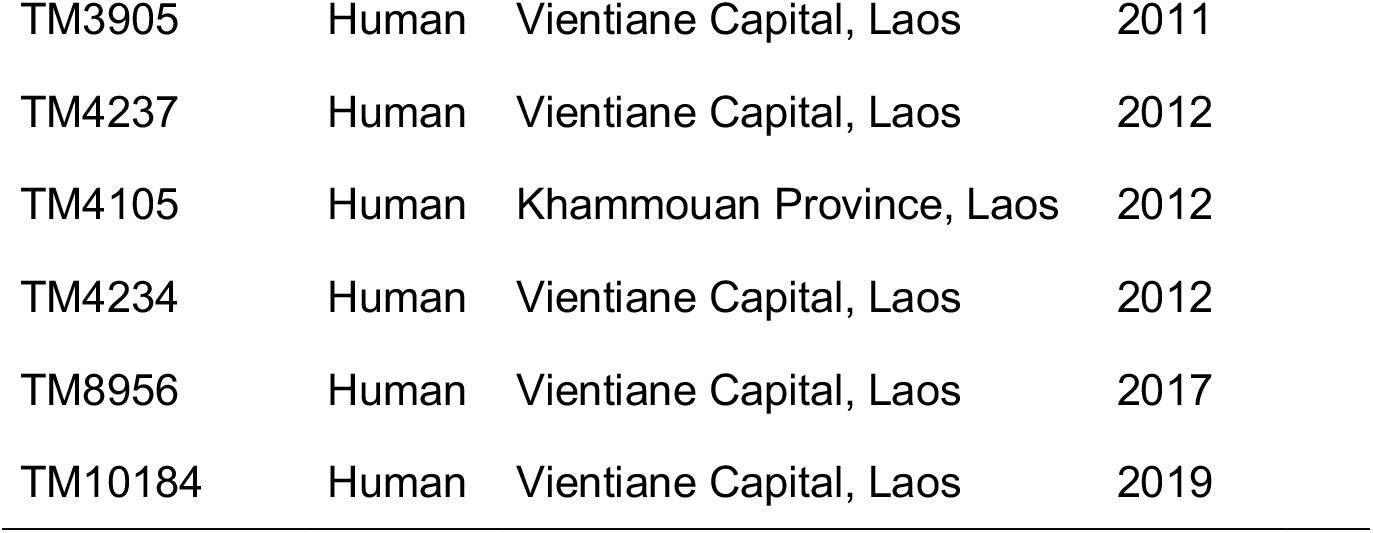
Isolate information table.

| **Isolates** | **Host** | **Country** | **Year** | **Reference** |
| --- | --- | --- | --- | --- |
| NC_006142.1 /Wilmington | Human | USA | 1928 | Maxcy, 1928 [31] |
| GEAR | Human | South Africa | 1939 | Gear, 1954 [32] |
| GER | Human | Azerbaijan | 1946 | Eremeeva et al, 1996 [33] |
| MUSSEIBOV | Human | Azerbaijan | 1949 | Eremeeva et al, 1996 [33] |
| FLA6590 | Rat | USA | 1951 | Perez Gallardo and Fox, 1948 [34] |
| TA837 | Rat | Thailand | 1963 | WRAIR, Bozeman/Sangkasuwan |
| TH1527 | Human | Thailand | 1965 | WRAIR, Bozeman/Sangkasuwan |
| PAKNA | Rat | Pakistan | 1970 | University of Maryland, Dasch/Wisseman |
| B9991CWPP | Rat | Myanmar | 1970 |  |
| AZ306 | Rat | Ethiopia | 1975 | University of Maryland, A. Azad, personal communication |
| AZ331 | Rat | Ethiopia | 1975 | University of Maryland, A. Azad, personal communication |
| TM1041 | Human | Vientiane Capital, Laos | 2006 |  |
| TM1377 | Human | Vientiane Capital, Laos | 2007 |  |
| TM2418 | Human | Vientiane Capital, Laos | 2008 |  |
| TM2522 | Human | Vientiane Capital, Laos | 2009 |  |
| TM2607 | Human | Vientiane Capital, Laos | 2009 |  |
| TM2529 | Human | Vientiane Capital, Laos | 2009 |  |
| TM2557 | Human | Vientiane Capital, Laos | 2009 |  |
| TM2540 | Human | Bolikhamxay Province, Laos | 2009 |  |
| TM2509 | Human | Vientiane Capital, Laos | 2009 |  |
| TM3627 | Human | Vientiane Capital, Laos | 2010 |  |
| TM4034 | Human | Vientiane Capital, Laos | 2011 |  |

|  |  |  |  |
| --- | --- | --- | --- |
| TM3905 | Human | Vientiane Capital, Laos | 2011 |
| TM4237 | Human | Vientiane Capital, Laos | 2012 |
| TM4105 | Human | Khammouan Province, Laos | 2012 |
| TM4234 | Human | Vientiane Capital, Laos | 2012 |
| TM8956 | Human | Vientiane Capital, Laos | 2017 |
| TM10184 | Human | Vientiane Capital, Laos | 2019 |

**Table 2.**
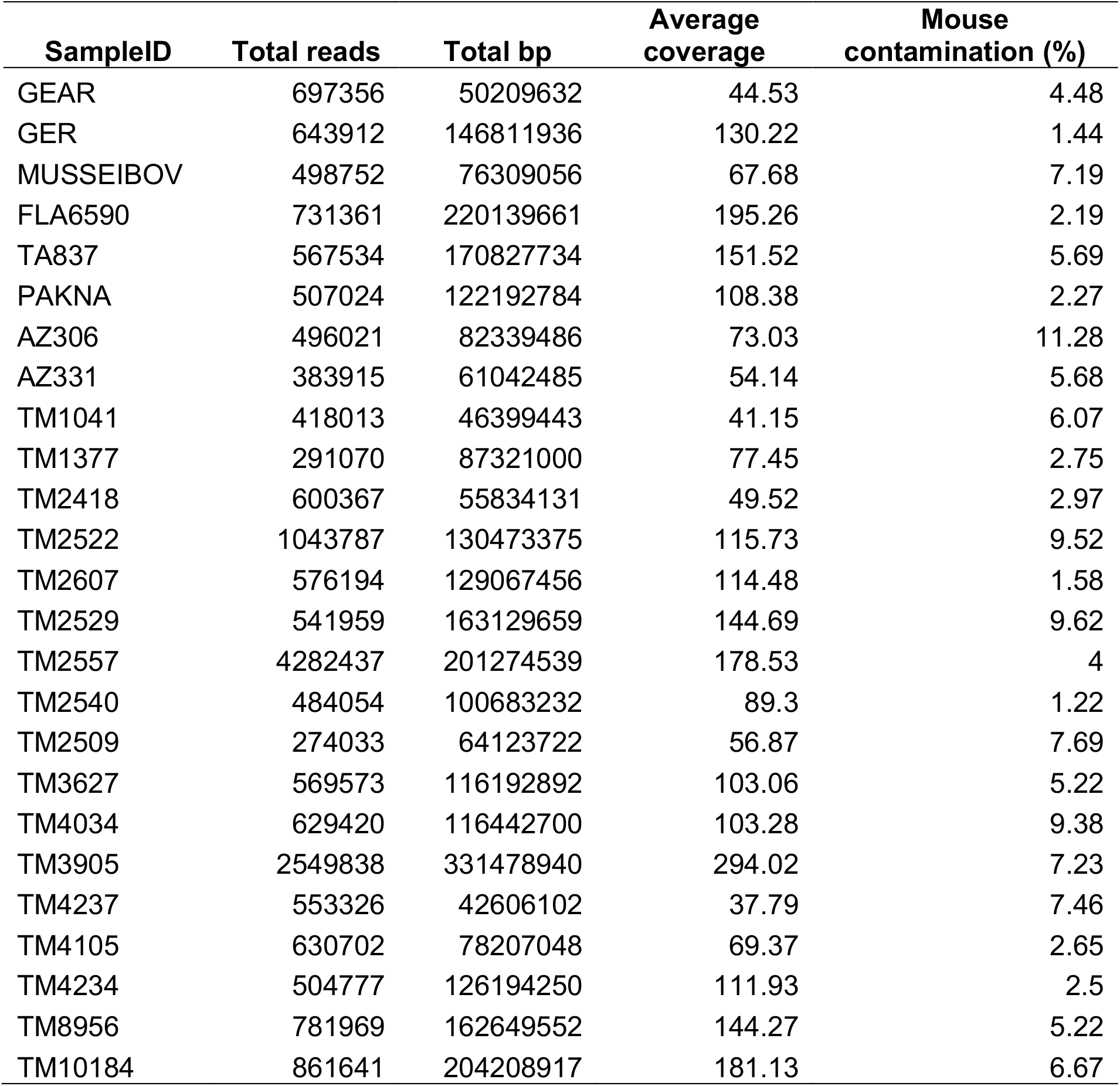
Paired-end sequencing data summary.

| **SampleID** | **Total reads** | **Total bp** | **Average coverage** | **Mouse contamination (%)** |
| --- | --- | --- | --- | --- |
| GEAR | 697356 | 50209632 | 44.53 | 4.48 |
| GER | 643912 | 146811936 | 130.22 | 1.44 |
| MUSSEIBOV | 498752 | 76309056 | 67.68 | 7.19 |
| FLA6590 | 731361 | 220139661 | 195.26 | 2.19 |
| TA837 | 567534 | 170827734 | 151.52 | 5.69 |
| PAKNA | 507024 | 122192784 | 108.38 | 2.27 |
| AZ306 | 496021 | 82339486 | 73.03 | 11.28 |
| AZ331 | 383915 | 61042485 | 54.14 | 5.68 |
| TM1041 | 418013 | 46399443 | 41.15 | 6.07 |
| TM1377 | 291070 | 87321000 | 77.45 | 2.75 |
| TM2418 | 600367 | 55834131 | 49.52 | 2.97 |
| TM2522 | 1043787 | 130473375 | 115.73 | 9.52 |
| TM2607 | 576194 | 129067456 | 114.48 | 1.58 |
| TM2529 | 541959 | 163129659 | 144.69 | 9.62 |
| TM2557 | 4282437 | 201274539 | 178.53 | 4 |
| TM2540 | 484054 | 100683232 | 89.3 | 1.22 |
| TM2509 | 274033 | 64123722 | 56.87 | 7.69 |
| TM3627 | 569573 | 116192892 | 103.06 | 5.22 |
| TM4034 | 629420 | 116442700 | 103.28 | 9.38 |
| TM3905 | 2549838 | 331478940 | 294.02 | 7.23 |
| TM4237 | 553326 | 42606102 | 37.79 | 7.46 |
| TM4105 | 630702 | 78207048 | 69.37 | 2.65 |
| TM4234 | 504777 | 126194250 | 111.93 | 2.5 |
| TM8956 | 781969 | 162649552 | 144.27 | 5.22 |
| TM10184 | 861641 | 204208917 | 181.13 | 6.67 |

### Variant calling and phylogeny

We identified 153 single nucleotide polymorphisms (SNPs) among the 28 strains of *R. typhi*. The majority of variants were annotated as missense variants compared to the Wilmington reference strain (Table 3). A phylogenetic tree generated from the 153 SNPs found in 28 *R. typhi* strains (Figure 1), shows that the isolates cluster into two distinct groups: the historic group from 1928 to 1975, and the modern group from 2006 to 2019. One isolate of the modern group, TM8956, was placed in the historic group instead of the modern group, and clusters most closely with two isolates from Thailand. The two isolates from Thailand (TH1527 and TA837) and Ethiopia (AZ330 and AZ306) form geographical clusters, while the other historic strains with the same geographic origin do not cluster together. Strains also do not cluster by their original host, with the rat samples separated across the tree, emphasizing the cross-species nature of this intracellular organism.

**Table 3.** Variant annotation of 27 *R. typhi* samples.

| Annotation | N | Percentage (%) |
| --- | --- | --- |
| Intergenic region | 38 | 24.84 |
| Missense variant | 78 | 50.98 |
| Stop retained variant | 1 | 0.65 |
| Synonymous variant | 36 | 23.53 |
| Total | 153 | 100 |

**Figure 1.**
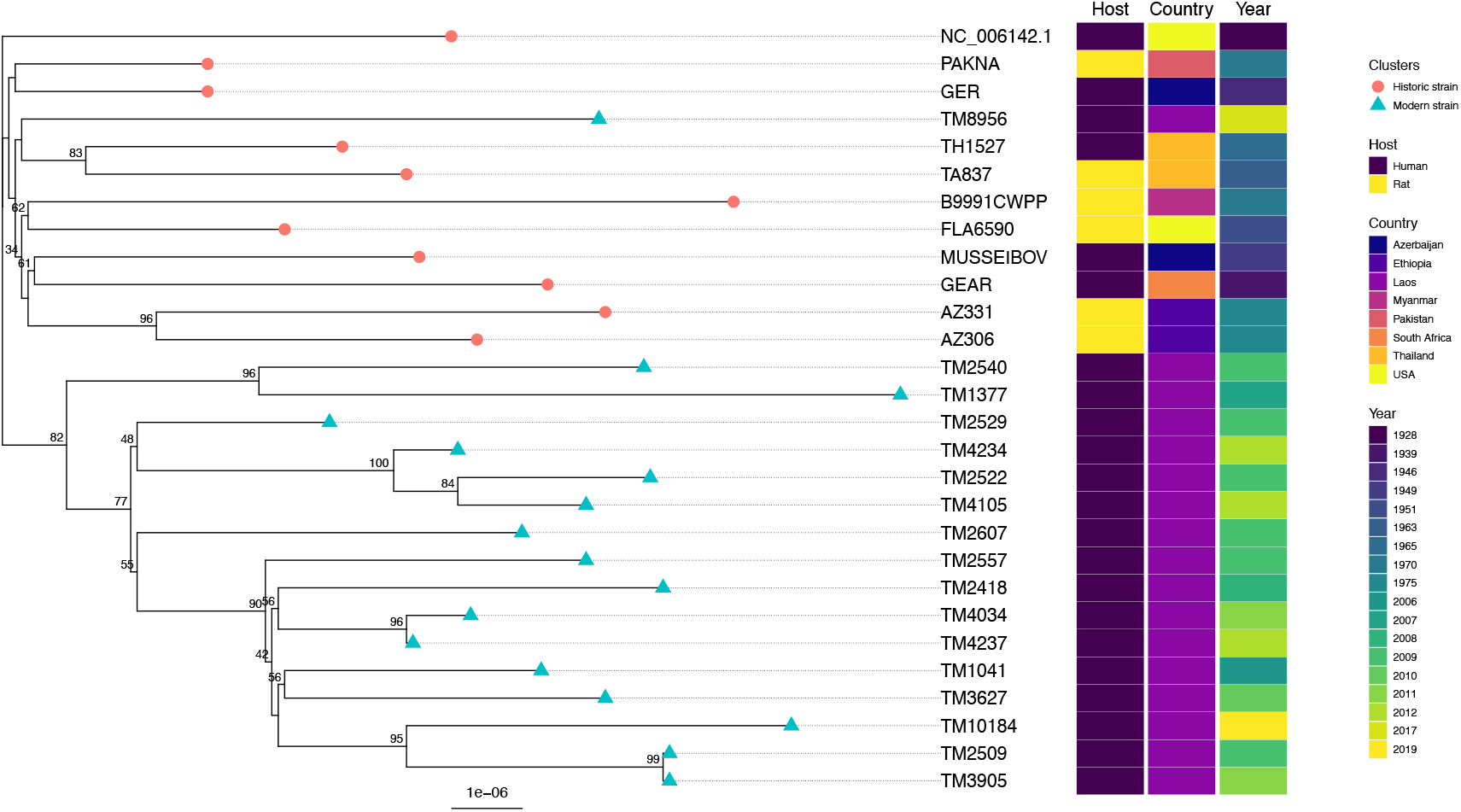
Phylogenetic tree of 28 *Rickettsia typhi* isolates. The phylogenetic tree was constructed using Iqtree with 1,000 bootstrap replicates. Figure indicates bootstrap support. The scale bar shows the substitution rate. The tree was plotted, annotated, and visualized by ggtree.

Kato *et. al*. [13] genotyped SNPs and indels identified using the three complete genomes in a wider set of strains, which includes the AZ306, AZ331, TA837, PAKNA (also called NA18), MUSSEIBOV, GER, GEAR, and FLA6590 strains which were also sequenced in our study. We compared our variant calls across 22 SNPs and two small indels (i3_162 and i10_825) identified by Kato *et al*. [13], and the calls were identical across the strains shared between the two studies. With the extra SNPs we identify in this study it is possible to distinguish the PAKNA and GER strains, which cannot be separated using the smaller Kato set of SNPs and indels.

We looked for clock signals in the mutation rate in *R. typhi* using BactDating which dates the nodes of a bacterial phylogenetic tree. Figure 2 shows the root-to-tip distance generated from the phylogenetic tree. Although there is a weak correlation between sampling date and root-to-tip distance, the signal is not significant after testing against a model with equal clock dates. Model comparison using Deviance Information Criterion (DIC) values showed a lower value for the clock model (DIC = 0.00) than the no-clock (equal-dates) model (DIC = 0.37), indicating that the clock model does not substantially improve the fit over the null model, suggesting insufficient evidence for a consistent molecular clock signal in this dataset.

**Figure 2.**
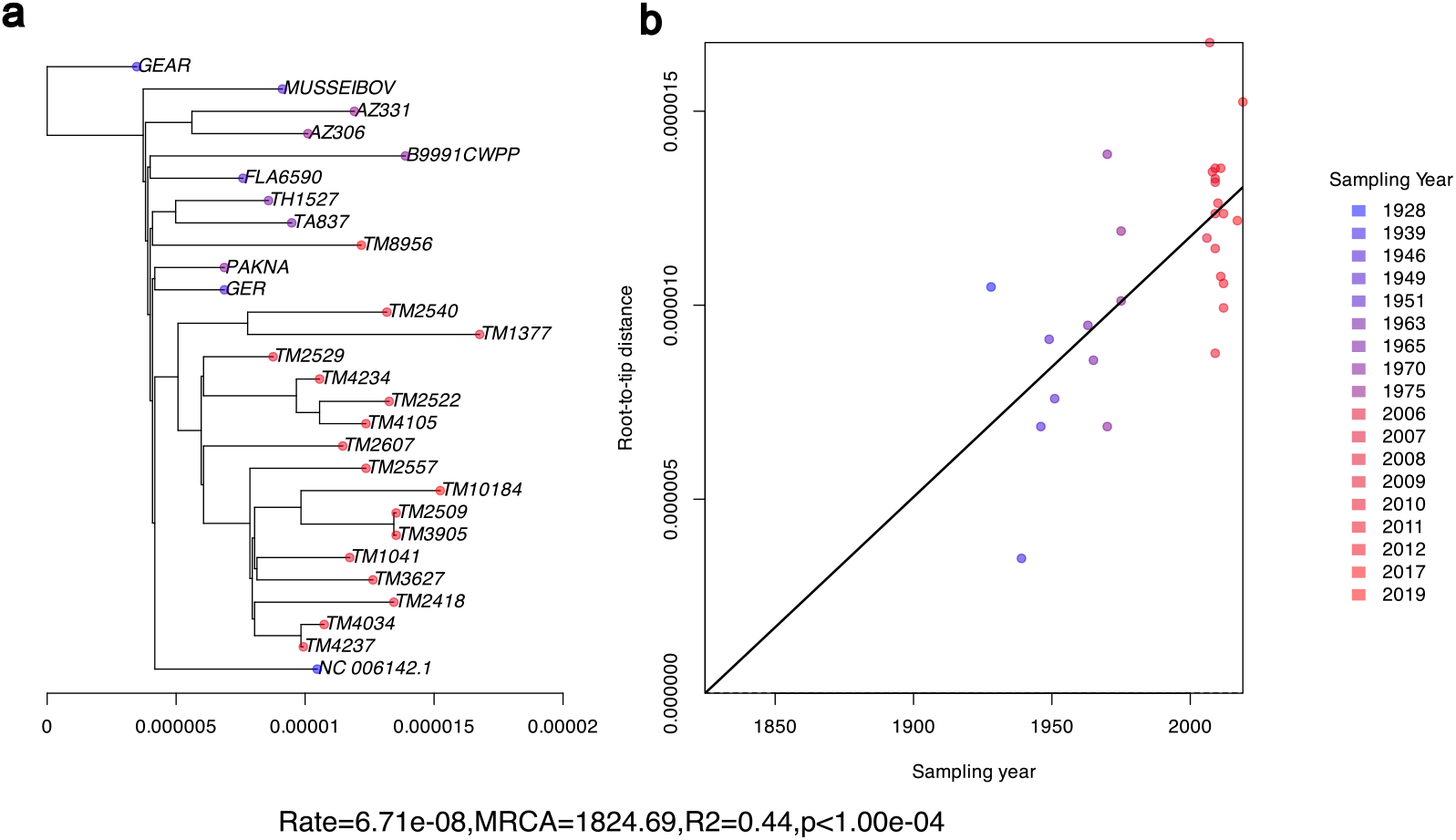
a) Phylogenetic tree generated using Iqtree showing substitution rate on the x-axis. Tree tips are coloured according to sampling date for each strain, with the blue to red gradient showing earliest to most recent strains. b) Root-to-tip regression showing a weak association between sampling date and root-to-tip distance.

Nucleotide diversity and selection pressures were estimated by SNPgenie. The results show that the overall nucleotide diversity (π) is low at 1.27e^−05^ overall, indicating limited genetic variation within the population. The nonsynonymous diversity (πN) of 1.14e^−05^ is slightly lower than synonymous diversity (πS) of 1.28e^−05^, suggesting purifying (negative) selection.

We identified one isolate, GEAR, with a nucleotide substitution in the *rpoB* gene, which encodes the RNA polymerase beta subunit and has previously been implicated in rifampin resistance. We observered an nonsynoymous A to C substitution at genomic position 164,354 resulting with the replacement of asparagine with histidine at residue 1039. This mutation does not correspond to the amino acid substitutions previously observed in the rifampin-resistant *rpoB* gene mutants generated in the laboratory [8]. However, the transition from asparagine to histidine introduces a larger side chain residue into *rpoB*.

### Assembly and genome annotation

The Illumina short-read FASTQ files were *de novo* assembled using Skesa and annotated using Prokka. Contigs which showed 99% or greater similarity to the mouse genome using BLAST were assumed to be culture contaminants and were removed. The assembly statistics for each isolate are shown in Table 4.

**Table 4.** Assembly and annotation statistics.

| **Isolate** | **Contigs** | **Size (bp)** | **Non-mouse contigs** | **Size(bp) after mouse contamination removal** | **N50** | **CDS** | **tRNA** | **rRNA units** |
| --- | --- | --- | --- | --- | --- | --- | --- | --- |
| TM4237 | 11 | 1109632 | 9 | 1108517 | 274951 | 861 | 33 | 3 |
| TM4105 | 10 | 1110196 | 8 | 1108685 | 273807 | 863 | 33 | 3 |
| TM4034 | 7 | 1108600 | 7 | 1108600 | 274951 | 863 | 33 | 3 |
| TM4234 | 6 | 1110100 | 6 | 1110100 | 656226 | 865 | 33 | 3 |
| TM2522 | 18 | 1118980 | 5 | 1109506 | 656775 | 864 | 33 | 3 |
| TM2607 | 6 | 1109456 | 6 | 1109456 | 274951 | 862 | 33 | 3 |
| TM2529 | 7 | 1126548 | 5 | 1109394 | 314979 | 864 | 33 | 3 |
| TM3627 | 14 | 1111656 | 10 | 1108893 | 248264 | 861 | 33 | 3 |
| TM3905 | 28 | 1135971 | 5 | 1109340 | 656202 | 863 | 33 | 3 |
| TM2557 | 14 | 1115564 | 7 | 1109043 | 274951 | 862 | 33 | 3 |
| TM8956 | 11 | 1111254 | 7 | 1108880 | 620152 | 861 | 33 | 3 |
| TM2540 | 9 | 1109880 | 8 | 1109298 | 160875 | 863 | 33 | 3 |
| TM10184 | 21 | 1128700 | 8 | 1109070 | 274951 | 862 | 33 | 3 |
| TM1041 | 14 | 1112308 | 10 | 1109445 | 187840 | 863 | 33 | 3 |
| AZ306 | 15 | 1130205 | 6 | 1109970 | 389013 | 863 | 33 | 3 |
| AZ331 | 9 | 1125476 | 7 | 1109262 | 274951 | 863 | 33 | 3 |
| FLA6590 | 10 | 1123985 | 6 | 1109047 | 574761 | 863 | 33 | 3 |
| GEAR | 14 | 1115426 | 9 | 1109679 | 274951 | 864 | 33 | 3 |
| GER | 16 | 1121520 | 11 | 1109339 | 220491 | 864 | 33 | 3 |
| PAKNA | 9 | 1126848 | 7 | 1109757 | 248253 | 862 | 33 | 3 |
| MUSSEIBOV | 9 | 1110795 | 7 | 1108930 | 274951 | 861 | 33 | 3 |
| TA837 | 10 | 1111092 | 9 | 1110293 | 275097 | 865 | 33 | 3 |
| TM2418 | 7 | 1111286 | 5 | 1109330 | 620201 | 864 | 33 | 3 |
| TM1377 | 11 | 1110450 | 10 | 1109441 | 275504 | 865 | 33 | 3 |
| TM2509 | 10 | 1110369 | 8 | 1109130 | 274918 | 862 | 33 | 3 |
| NC_006142.1 | 1 | 1111496 | 1 | 1111496 | 1111496 | 865 | 33 | 3 |
| TH1527 | 1 | 1112372 | 1 | 1112372 | 1112372 | 864 | 33 | 3 |
| B9991CWPP | 1 | 1112957 | 1 | 1112957 | 1112957 | 864 | 33 | 3 |

The gene content was assessed using BUSCO (Figure S1). The number of complete genes in the assemblies is the same as for the Wilmington, TH1527, and B9991CWPP reference: 86.3%, 0.8%, and 12.9% for complete, fragmented, and missing genes, respectively, indicating that our assemblies are likely complete in gene content in comparison to these complete genome assemblies.

Using Quast to compare our assembled contigs to the Wilmington reference, we identify two regions with potential assembly errors. These are around positions 185,255bp to 321,747bp and 841,954bp to 907,053bp, where there is a discontinuity in the alignment of the assembled contigs to the reference genome (Supplementary Figure 2), likely caused by difficulty in resolving the genome sequence around these regions during the genome assembly process.

### Pan-genome

We constructed a pangenome from all of the strains described in this paper using Panaroo (Table 4), showing 861 core genes and 6 accessory genes across the 28 strains. 861-865 coding sequences were annotated across the strains, suggesting the majority of genes in *R. typhi* are present in the core genome. Six accessory gene groups were present in only a subset of strains, and we investigated in detail as these represent the only potential non-core genes in the *R. typhi* genome (Supplementary Information). By aligning sequences from these gene groups to the genome assemblies and identifying the region where genes were missing from assemblies using synteny, we determined that these six groups did not represent real accessory genes which are differentially present in *R. typhi*, but were artifacts of genome assembly (Supplementary Figures S3–S9).

### Resequencing of TM2540

TM2540 was previously sequenced and assembled using a combination of Oxford Nanopore MinION and Illumina MiSeq sequencing, with the Oxford Nanopore long reads used to produce a draft assembly which was polished using Illumina data [12]. The work by Kato *et. al*. suggested that this isolate had an unusually high number of variants compared to other strains, therefore we resequenced this isolate using Illumina technology to identify whether the original assembly had spurious SNP calls. We compared the variant calls generated from Snippy using the previous assembly and the newest Illumina MiSeq data and found that the number of SNPs identified using our newer Illumina dataset only is the same as previously described, with 17 SNPs in comparison to other strains. These 17 SNPs are not identified as errors by polishing the genome with Pilon using the newest Illumina data, and TM2540 is positioned with the other modern Laos isolates in our phylogeny. However, when we compare indel calls from Snippy using the new Illumina data and the original assembly, we called three indels using the new Illumina data compared to 20 in the old assembly. After polishing the original assembly with the new Illumina data, we removed 12 of the 20 indels, indicating potential excess indel calls when using the previously published Oxford Nanopore data for assembly.

## Discussion

We demonstrate that the mutation rate across *R. typhi* is extremely low, consistent with previous work [12,13], and the pangenome is almost non-existent with very little accessory genome present across the strains tested. This is consistent with the biology of *R. typhi*, an obligate intracellular bacterium with a reduced genome that typically experiences limited exposure to mutagenic environmental factors and horizontal gene transfer [6]. Recent work has confirmed that lower pangenome fluidity is correlated with the obligate intracellular lifestyle, and is likely due to lifestyle factors and not effective population size, which is also smaller in intracellular parasites [35]. *Rickettsia* genomes show substantial gene loss, common among intracellular organisms that rely heavily on their host for nutrition and other metabolic processes [36]. The lack of genome fluidity in *R. typhi* suggests all inessential genes have been lost from the genome, and that either gene gain via horizontal transfer from other species occurs rarely in *R. typhi*, or selective pressure leads to loss of any inessential genes gained.

Despite the low number of mutations seen across this species, we identified some temporal clustering in the strains examined, with two broad clades representing the historic and modern strains. As all the modern strains in this study were collected from patients in Laos, it is difficult to determine whether the same temporal clustering would be observed with a more geographically diverse data set of modern strains. Notably, there was a single Laos strain which clustered with the historic strains which was most closely related to two historic strains collected in Thailand. It is not possible to determine whether these historic clade strains were continually circulating in Laos, or represents a re-introduction of these strains, potentially from Thailand as closely related strains were demonstrated to be circulating in Thailand in the past. The historical strains used in this study will have undergone many passages in culture, while the modern strains from Laos are low-passage strains sampled as close to possible to the date when the patient presented with *R. typhi* disease. The clustering of a modern Lao strain with the historical strains suggests that the variants seen are circulating in patient strains, and do not represent variation which has arisen through adaptation to culture.

Similarly, our modern Lao strains all have a human origin, but among the historical strains there is no clustering by species of origin, suggesting that *R. typhi* strains do not show distinct genotypes adapted to different mammalian hosts. A weakness of this study is that there are no available genomes from strains obtained directly from fleas, the main vector of *R. typhi*, and it is possible that the genetic diversity of strains in the vector is greater than those which can infect a mammalian host.

We detect one isolate, collected in 1939 in South Africa, with a mutation in the *rpoB* gene, which has previously been implicated in laboratory-induced rifampicin resistance. The mutation we found has not been previously reported to cause rifampicin resistance, and it is unknown whether this mutation contributes to phenotypic resistance in this strain, which was collected before rifampicin was in use. A study comparing the sequences of naturally rifampin-resistance *Rickettsia* to sensitive species hypothesized that specific pairs of residues introducing larger side chain residues could result in reduced binding efficacy for rifampin and lead to resistance [10]. The mutation in the GEAR isolate introduces a larger side chain residue which could lead to steric hindrance of the rifampin binding site. However, the mutations at positions 409 and 973, which are present in all naturally resistant *Rickettsia*, were not seen in any of our isolates.

In this study we used Illumina short-read sequence data to assemble the new strains. Despite the short read length, it was possible to assemble each genome in 11 or fewer contigs, and with only two regions showing possible misassemblies, demonstrating that while long-read sequencing is beneficial for genome assembly it is possible to obtain high-quality genomes with short-read sequencing alone. A target capture sequencing approach using probes designed to hybridise to reference genomes works effectively for *O. tsutsugamushi* [37] and *R. prowazekii* [38], and given the small genome size and limited diversity of *R. typhi* a similar approach could be used to sequence genomes from clinical specimens and potentially from vector hosts. As part of this study, we re-sequenced the TM2540 strain, which has an existing complete genome generated using a hybrid assembly of older Oxford Nanopore and Illumina data. The SNP calls generated from the resequenced strain matched the original assembly, but some of the indels previously identified using Oxford Nanopore data could not be validated using the newer data, suggesting they were artifacts remaining from the assembly process. The previous assembly was generated using R9.4.1 flow cells, which is known to give data with a higher error rate than the newer and more accurate R10 flow cells and kits [39].

The extremely low genetic diversity of *R. typhi* across isolates collected almost a century apart from different continents has implications for investigation into *R. typhi* pathogenesis, diagnostics, treatment, and prevention. Severity in *R. typhi* disease is unlikely to have a pathogen genetic component, and diagnostics which rely on nucleic acid amplification are unlikely to be affected by sequence variation. While rifampicin resistance has been seen in a laboratory strain of *R. typhi* [8], naturally-occurring resistance will need to overcome the low mutation rate and strong selective pressure and, to our knowledge, has not yet arisen in Laos PDR. The lack of variation may also be beneficial in development of *R. typhi* vaccines and point-of-care diagnostic tools, as there is minimal variation in possible antigenic targets.

## Acknowledgements

This work was supported in part by the Wellcome Trust (grant 220211/Z/20/Z). For the purpose of Open Access, the author has applied a CC BY public copyright license to any Author Accepted Manuscript version arising from this submission. The funding source had no involvement in the design, methods or analysis of the study. We are grateful to the director of Mahosot Hospital, the director and staff of the Microbiology Laboratory, and the ward doctors and nurses at Mahosot Hospital and the late Dr Rattanaphone Phetsouvanh for her leadership and research on rickettsial pathogens. We acknowledge Professor Sabine Dittrich, Phonepasith Panyanouvong, Davanh Sengdatka, Thaksinaporn Thaojaikong and Narongchai Tongyoo who grew the initial patient isolates used for this study and supervised the laboratory.

## Ethical statement

Sequencing was carried out from strains previously isolated from patient specimens during routine microbiological investigations. Strains were grown in commercial cell lines and no sequencing was performed from direct patient samples, therefore ethical approval for sequencing of isolates was not required as no patient sample or patient DNA was present.

## Supplementary data

**Figure S1.**
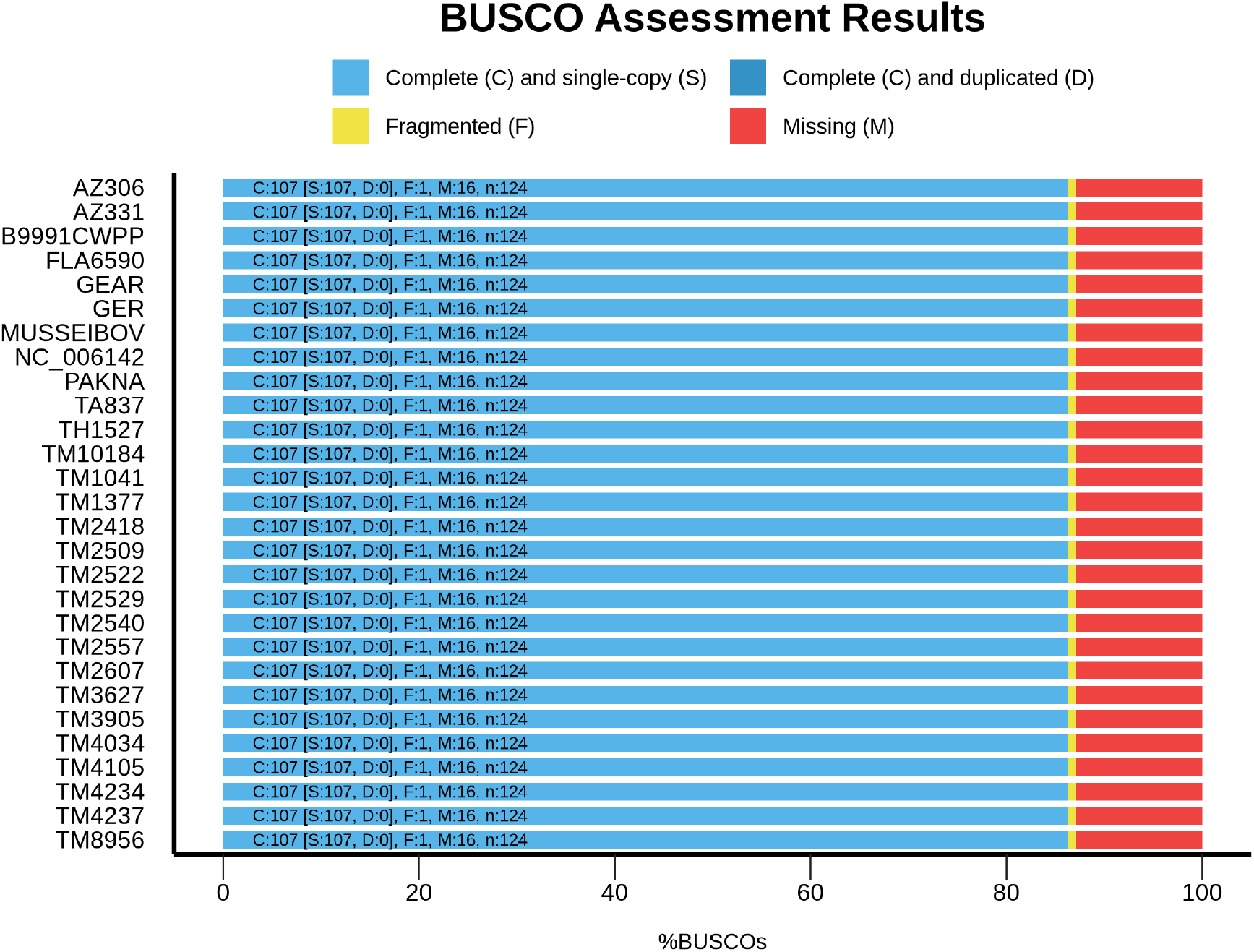
A quantitative assessment of the completeness in terms of expected gene content of a genome assembly by BUSCO, showing that the number of complete, fragmented, and missing genes is the same in the complete reference assemblies and our novel assemblies.

**Figure S2.**
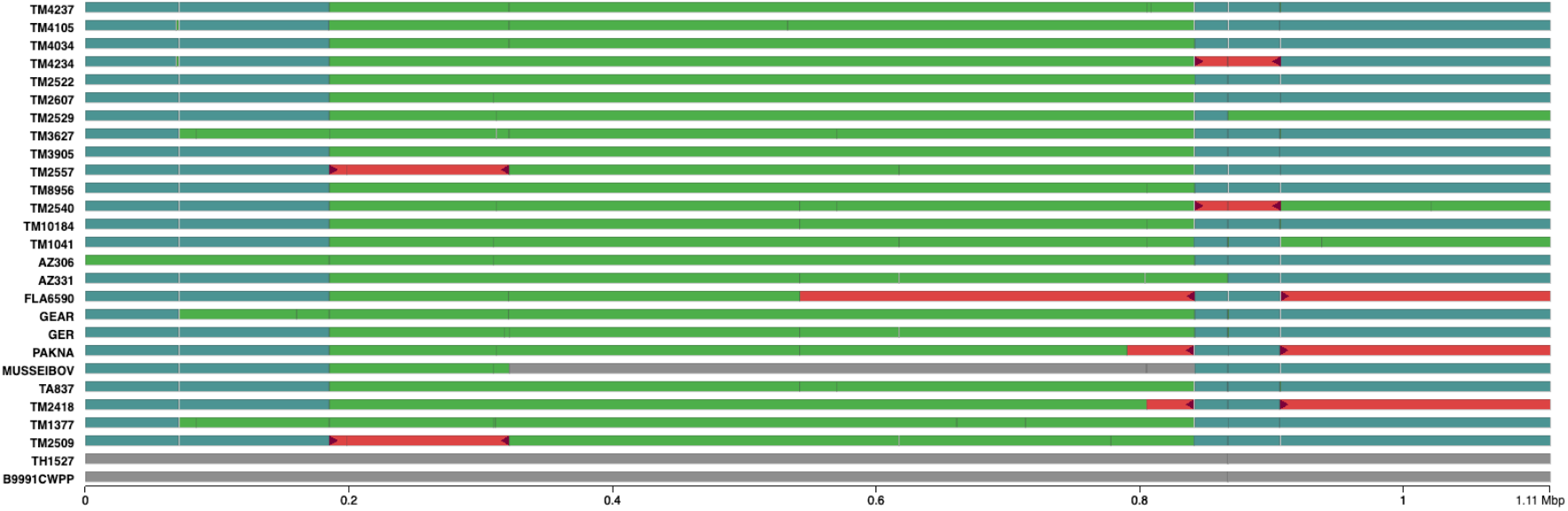
Contig alignment results to Wilmington strain by Quast and the Icarus contig viewer. The x-axis represents genome location in the Wilmington genome. Green and blue blocks represent correctly assembled contigs. Red blocks represent misassembled contigs which do not correspond to a single alignment block.

### 1. Investigation of accessory genes

Six gene groups were identified using Panaroo as potential accessory genes, present in less than 100% of the genomes. We investigated each gene group to determine if the gene was really not present in some strains or was missed due to errors in assembly or annotation of the genome, and concluded the genes were missed due to assembly and annotation errors..

#### a. Group_506

Group_506 is a hypothetical protein that is only present in the reference genomes and not the newly assembled strains (Figure S3). When the sequence of the gene from group_506 in genome NC_006142.1, B9991CWPP, and TH1527 were aligned to all genomes, it was present in all the genomes but with a similar sequence to genes annotated as group_505. Group_505 and group_506 are identical in length, 246bp, and have identical sequences (Figure S4). Group_506 is likely not present in the Illumina assembly data due to the difficulty of assembling very short duplicated regions using short-read sequencing technology. Using the Wilmington strain as a reference, the flanking genes between group_506 are the *tgt* gene and a hypothetical protein. We found that group_506 is missing from the short read assembly genomes and *tgt* is the last gene located at the end of a contig, indicating that group_506 cannot be assembled (Figure S5).

**Figure S3.**
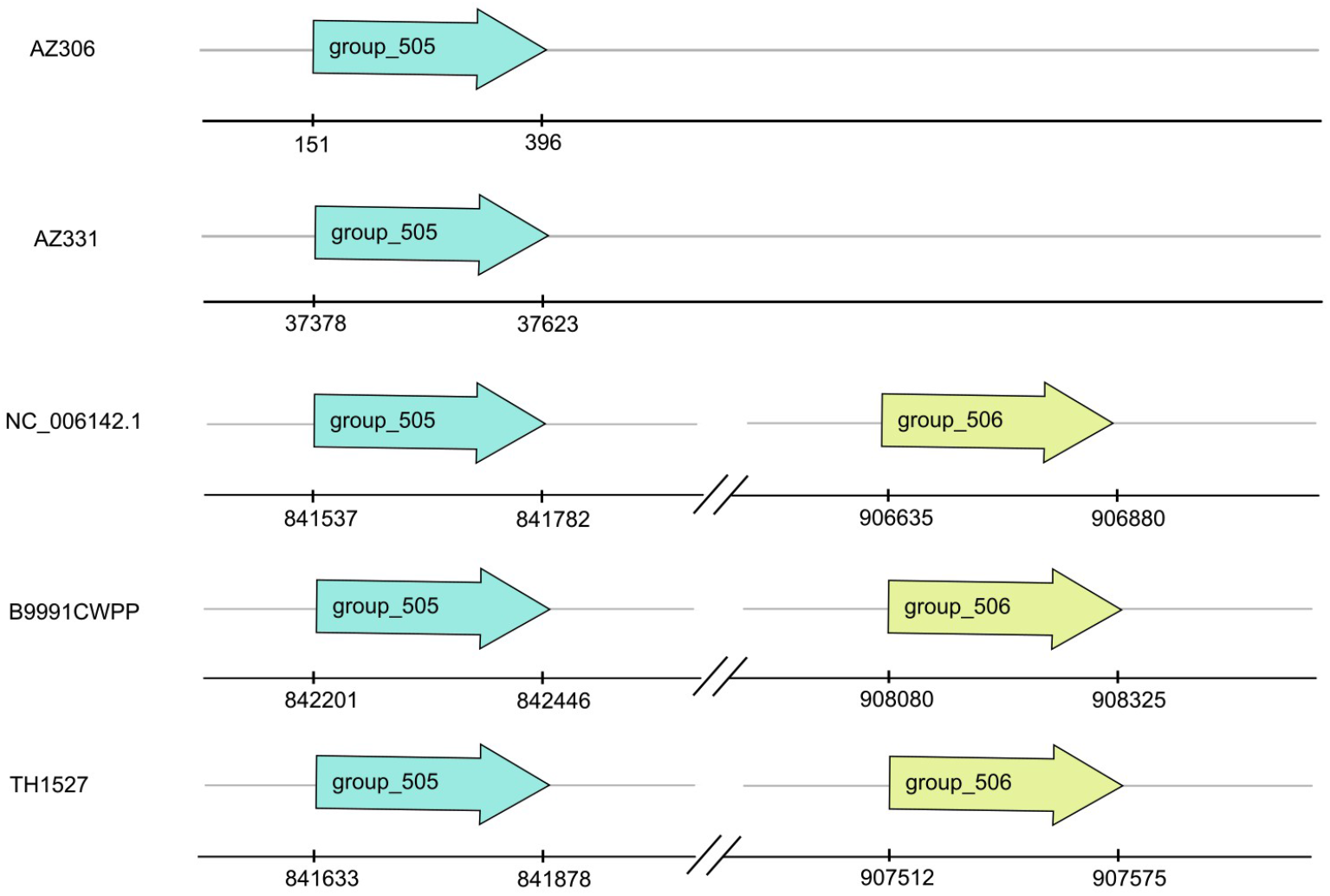
The illustration of group_506 and group_505 in the reference genomes (Wilmington, B991CWPP, and TH1527) and two selected short-read assembly genomes (AZ306 and AZ331). Positions give the location of this gene in the contig or genome.

**Figure S4.**
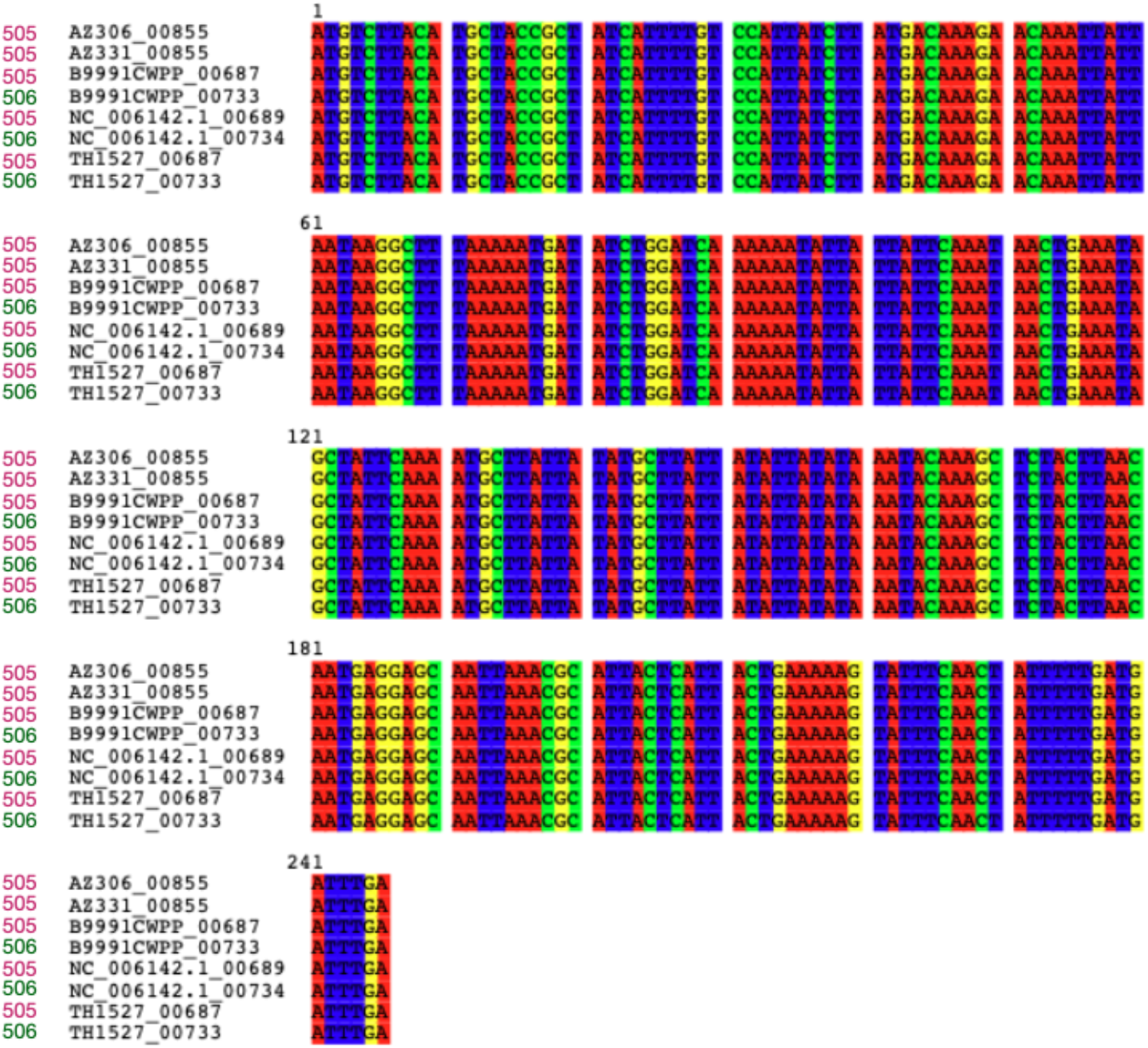
Multiple sequence alignment of group_505 and group_506 sequences.

**Figure S5.**
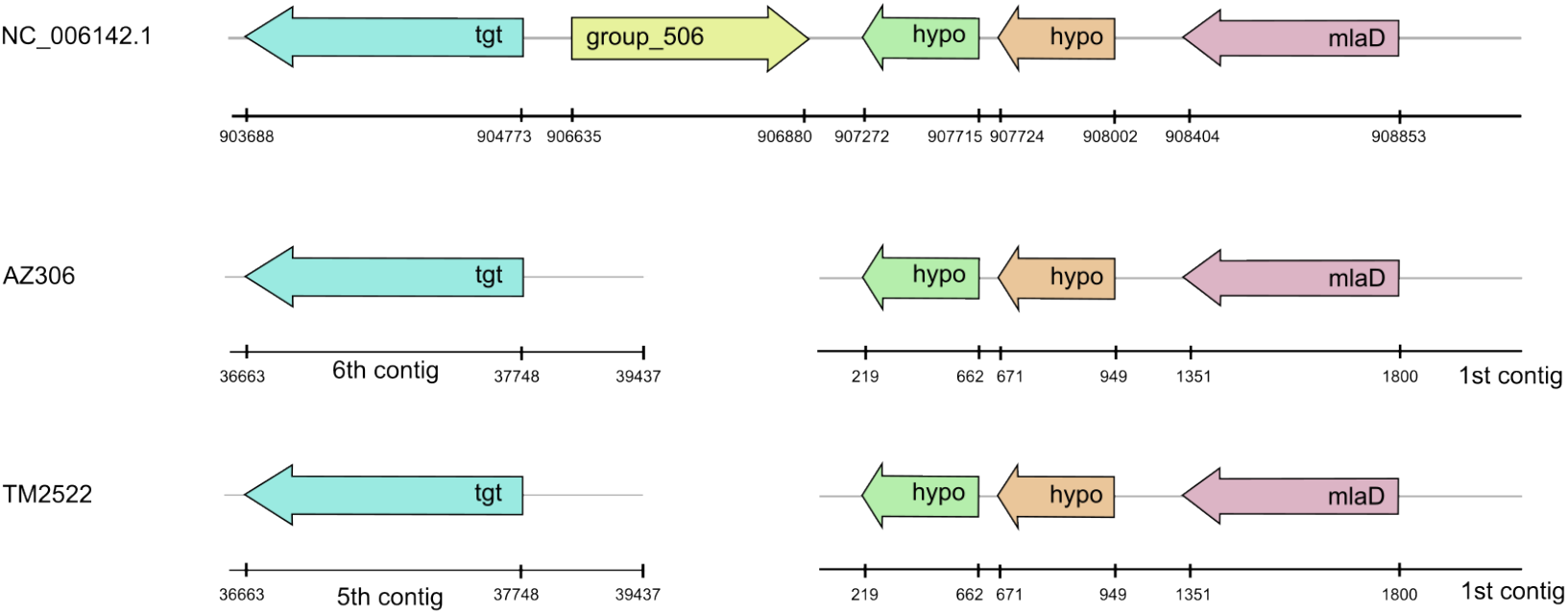
group_506 is absent in short read assembly genomes, AZ306 and TM2522.

#### b. Group_392

Group_392 is a hypothetical protein that is present in 24 strains, all of which were sequenced by Illumina. The coding sequence of group_392 was present in all the genomes, however in the samples where group_392 was not identified, the sequence was found to form part of the coding sequence of group_524 (Figure S6). A multiple sequence alignment of group_392 and group_524 shows that group_392 is a shortened fragment of group_524 due to this region being assembled as two separate contigs.

**Figure S6.**
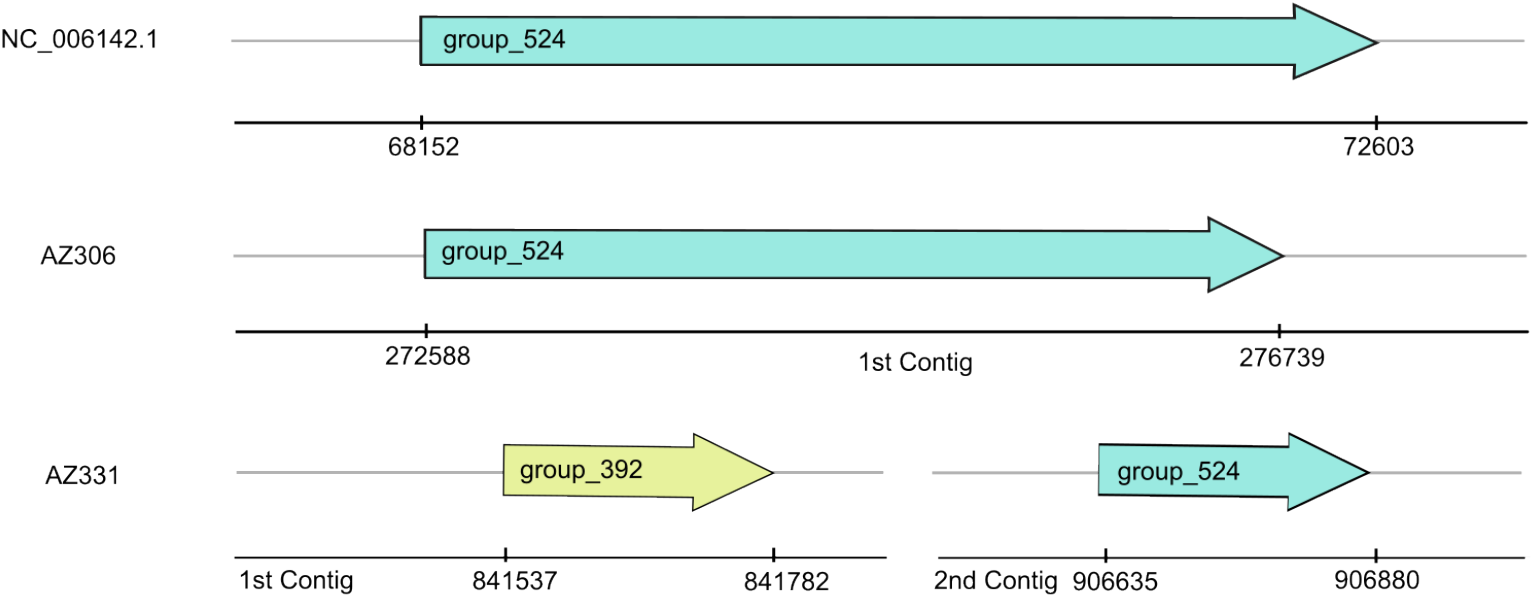
The illustration of group_392 and group_524 in assembled genome (NC_006142.1) and short-read assembly genomes (AZ306 and AZ331)

#### c. Group_509

Group_509 is a bifunctional (p)ppGpp synthetase/guanosine-3’5’-bis(diphosphate) 3’-pyrophosphohydrolase protein. The DNA sequence of the gene from group_509 was present in strain TM2607 but this strain did not have a gene present in group_509 as it is not annotated as having a valid coding sequence. Group_509 lies between group_260 and *groL* genes, and we selected DNA sequences from the end of group_260 to the start of *groL* genes of every absent sample and aligned with group_509 of the reference Wilmington strain. We found that the sequence of group_509 is there, but the contigs are fragmented in strains TM10184 and TM1041, and there is a sequence insertion between two parts of gene in the strains MUSSEIBOV, TM2607, and TM3627 (Figure S7).

**Figure S7.**
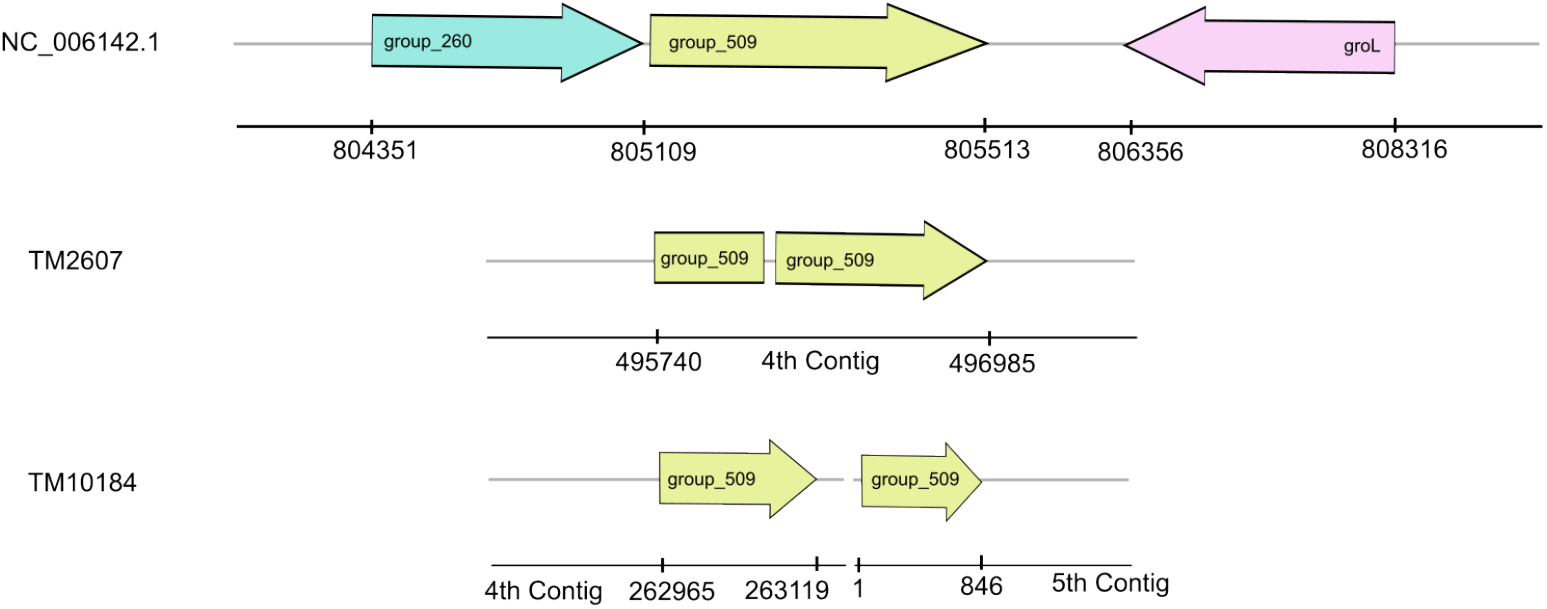
The illustration of group_509 in assembly genome (NC_006142.1) and short-read assembly genomes (TM2607 and TM10184)

#### d. Group_507, group_525, and group_512

Group_507 and group_525 are LicD family proteins, and group_512 is a phosphoryl choline transferase LicD protein. Group_507 and group_525 are never found in same strain. In strains where group_525 were found, it is always at the start of the contig, and the comparison of group_507 and group_525 by multiple sequence alignment has shown that group_525 is a shorter, truncated version of group_507; therefore, it is not clustered with group_507. TM2540 assembly data by Illumina lacked group_507, whereas in the TM2540 assembly data by Oxford Nanopore Technology (ONT) group_507 was present. From the MSA of group_507 and group_512 based on the assembled genomes, the length of both genes is nearly equal, around 800 base pairs, the sequences are very similar, and they are next to each other in the genome. The missing gene is likely due to the difficulties of assembling this tandem repeat gene.

**Figure S8.**
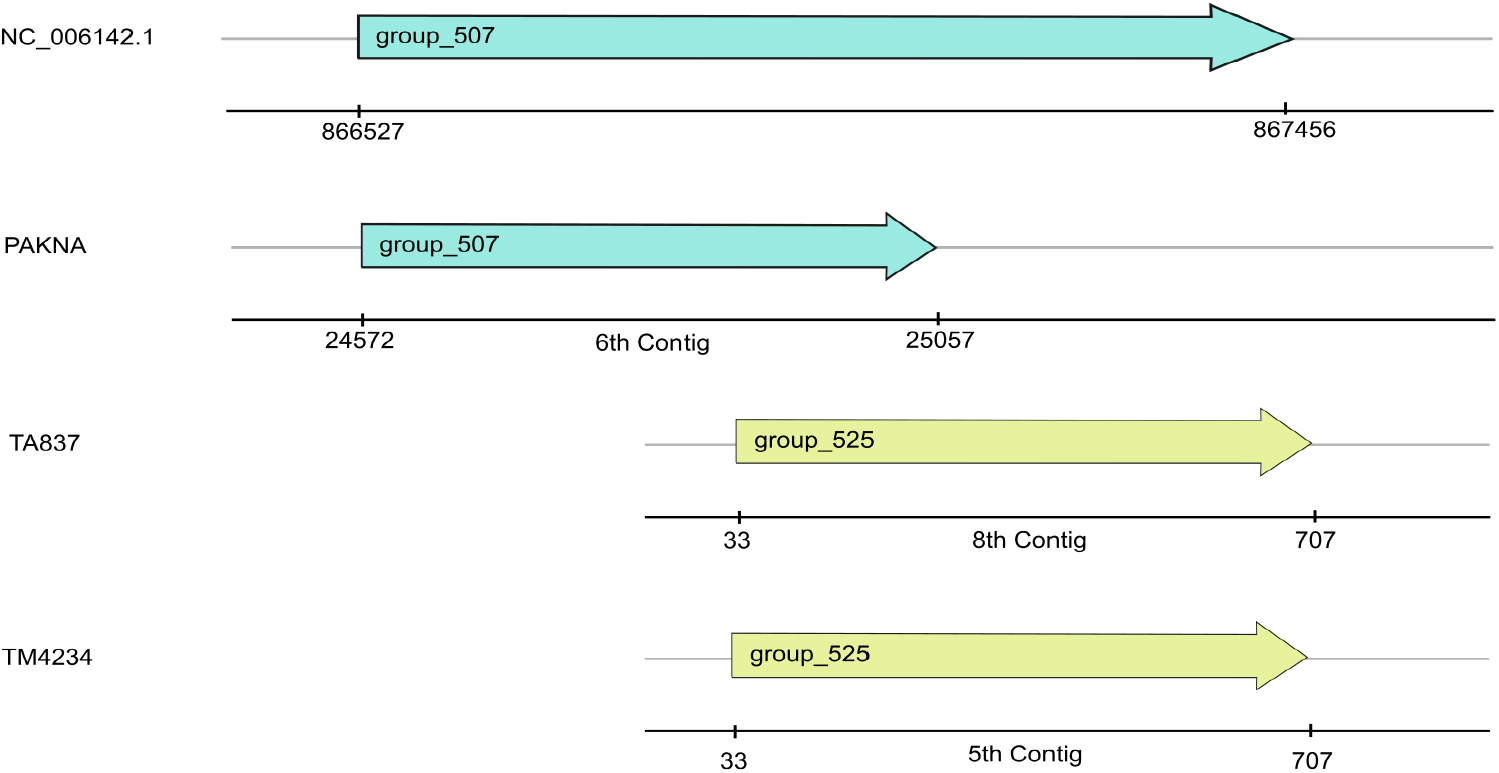
The illustration of group_507 and group_525 of assembly genome (NC_006142.1) and short-read assembly genomes (PAKNA, TA837, and TM4234)

**Figure S9.**
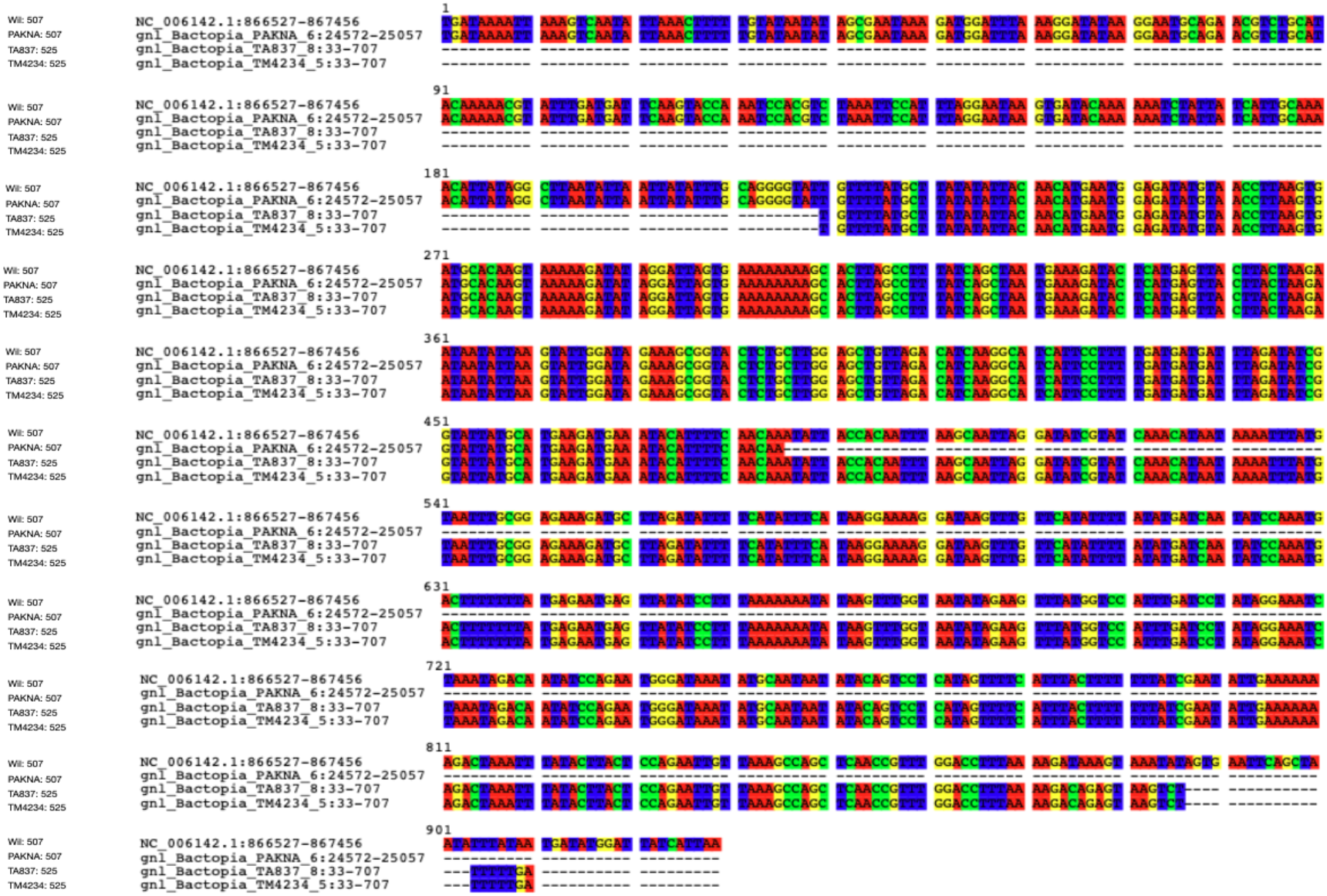
MSA of group_507 and group_525.

